# MADDD-seq, a novel massively parallel sequencing tool for simultaneous detection of DNA damage and mutations

**DOI:** 10.1101/2023.08.27.555013

**Authors:** Marc Vermulst, Samantha L. Paskvan, Claire Chung, Kathryn Franke, Nigel Clegg, Jennifer Madeoy, Annalyssa S. Long, Jean-Francois Gout, Jason H. Bielas

**Affiliations:** University of Southern California, Leonard Davis School of Gerontology, Los Angeles, CA; Translational Research Program, Public Health Sciences Division, Fred Hutchinson Cancer Research Center, Seattle, WA, USA; Department of Biological Sciences, Mississippi State University, Mississippi State, MS 39762, USA; Department of Laboratory Medicine and Pathology, University of Washington, Seattle, WA, USA

## Abstract

Our genome is exposed to a wide variety of DNA-damaging agents. If left unrepaired, this damage can be fixed into mutations that promote carcinogenesis and the development of genetically inherited diseases. As a result, it is crucial that we can detect DNA damage and mutations with exquisite sensitivity. Here, we describe a modified version of double barcoding sequencing technology termed Mutation And DNA Damage Detection-seq (MADDD-seq) that can detect DNA damage and mutations simultaneously, with a single assay. To demonstrate the utility of MADDD-seq as a multifunctional detection tool, we treated yeast cells with a DNA-damaging agent and tracked the presence of DNA damage and mutations over a 24-hour timespan. These experiments allowed us to identify thousands of adducts and mutations in a single sequencing run and expose the kinetics of DNA repair and mutagenesis in remarkable detail.

## INTRODUCTION

Our genome is exposed to a wide variety of DNA damaging agents, including radiation, metabolic side products, inflammatory molecules and viral particles(*1, 2*). This damage plays an important role in human pathology by encouraging apoptosis, promoting senescence, and accelerating the pace of human aging(*3*). In addition, DNA damage is highly mutagenic, and the mutations it creates can drive the evolution of human tumors(*4*). Accordingly, powerful tools are needed to detect DNA damage and mutations in human cells to advance our understanding of mutagenesis and inform strategies aimed at the prevention and treatment of disease.

Although various tools have been designed to detect bulky DNA adducts with high sensitivity(*5-9*), it has proven more difficult to detect adducts that are smaller in size(*10*). Small DNA adducts are important targets for biomedical research as well though, because they are more common than bulky adducts, and tend to be more amenable to translesion synthesis and thus mutation fixation(*10*). Accordingly, improved detection assays for small DNA lesions are urgently needed. A second limitation of current detection assays is that DNA damage and mutations are usually detected separately, with different assays. As a result, samples need to be split up into two different aliquots, each of which is manipulated in a unique way to accommodate the appropriate instrumentation. Due to these chemical and biological manipulations though, the samples are no longer identical, which could confound the experiment. This issue is further compounded by the use of two distinct detection instruments, whose sensitivity and specificity are never identical(*11, 12*). As a result, it is difficult to compare the readouts of DNA damage and mutation detection assays to each other in a truly quantitative fashion. Finally, the need for two separate assays also poses practical limitations for researchers, who will need to spend time and resources to perform their experiments.

To address these issues, we developed MADDD-seq, a novel massively parallel sequencing tool that can detect mutations and DNA adducts simultaneously, using a single assay and a single sample. Because MADDD-seq is a DNA-based sequencing tool, it can be applied to any organism of choice, and both damage and mutations can be mapped directly onto the genome. To do so, MADDD-seq advances on an extremely sensitive double barcoding strategy that we previously employed to detect mutations with unprecedented sensitivity(*13*). We then coupled these advances to a novel bio-informatic pipeline that uses the information encoded by our modified barcodes to identify mutations and DNA adducts in the same dataset and localize these adducts to a single base on a single strand of DNA. Because MADDD-seq depends on error-prone translesion synthesis during library amplification for the detection of DNA damage, we expect that MADDD-seq will be especially useful for the detection of small, highly mutagenic DNA adducts that drive mutagenesis in the aging process and various other forms of pathology.

To demonstrate the capabilities of MADDD-seq, and to interrogate the role of proliferation in mutagenesis, we treated dividing and arrested yeast cells with the alkylating agent methylnitronitrosoguanidine (MNNG), a chemical that acts primarily by adding alkyl groups to the O^6^ of guanine and O^4^ of thymine. Of these two lesions, O^6^-methyl guanine (O^6^-me-G) adducts have proven to be highly mutagenic(*14*). We then tracked these cells over a 24-hour timespan to determine how these lesions were distributed across the genome after exposure, how rapidly they were repaired, and how frequently they were fixed into mutations. Thus, MADDD-seq allowed us to monitor every step of the mutagenic process over time with a single assay to provide precise quantitative insight into the parameters that control our risk for disease.

## CONCEPT

Massively parallel sequencing tools have revolutionized modern medicine(*15*). They exposed the genetic heterogeneity of cancers, identified germline mutations responsible for human diseases, and opened the door for personalized medicine(*16*). After this initial wave of discoveries, researchers started to adapt massively parallel sequencing tools to detect mutations that are exceedingly rare, because they are only present in one, or a small number of cells in a patient(*11*). Although they are rare, random mutations hold incredible value for medical research. For example, by analyzing the spectrum of random mutations, the origins of human cancers can be exposed(*17*), while similar datasets can determine the impact of environmental mutagens on the human genome(*18*).

To do so, researchers had to modify massively parallel sequencing tools in various ways. The main issue they had to solve was that traditional sequencing data is rife with PCR artifacts and sequencing errors that are indistinguishable from *bona fide* mutations(*19, 20*). For example, DNA damage frequently introduces *ex vivo* mutations into sequencing libraries through error-prone translesion synthesis during construction steps that require PCR. To distinguish between true mutations and *ex vivo* mutations, several double-stranded barcoding strategies have been designed that tag both strands of a DNA duplex with a specific barcode(*21*). Accordingly, these techniques are usually referred to as “duplex-sequencing techniques”. The rationale behind duplex-sequencing is that if a mutation is present in a DNA molecule, it should be present in both of strands of the DNA duplex, while an artifact (such as an *ex vivo* mutation induced by DNA damage) should be present in only one(*21*). Because duplex sequencing allows the original DNA duplex to be reconstructed from sequencing data that covers both strands, it can error-correct massively parallel sequencing data.

The concept behind MADDD-seq is to take advantage of the process by which DNA damage can introduce *ex vivo* mutations into massively parallel sequencing libraries via PCR. We reasoned that the PCR steps of library construction techniques can only introduce mutations into early copies that are made of the damaged strand, while copies of the undamaged strand would remain artifact free. If so, it may be possible to use *ex vivo* mutations as reporters for the presence of DNA damage, by tracking them back to the original DNA strand from which they were derived. To do so, we biochemically designed our amplification reactions for efficient, error-prone bypass of DNA damage, and constructed a modified set of double-stranded barcodes that allows us to distinguish between copies that are made from the top, or the bottom strand of a DNA duplex (**fig. 1a**) with a custom-made bio-informatic pipeline that keeps track of *ex vivo* mutations s that arise in these copies (**fig. 1b**). Importantly, this strategy leaves the mutation detection component of the original “duplex sequencing” technology intact, so that MADDD-seq can detect both mutations and DNA damage simultaneously, with a single assay in a single sample.

**Figure 1.**
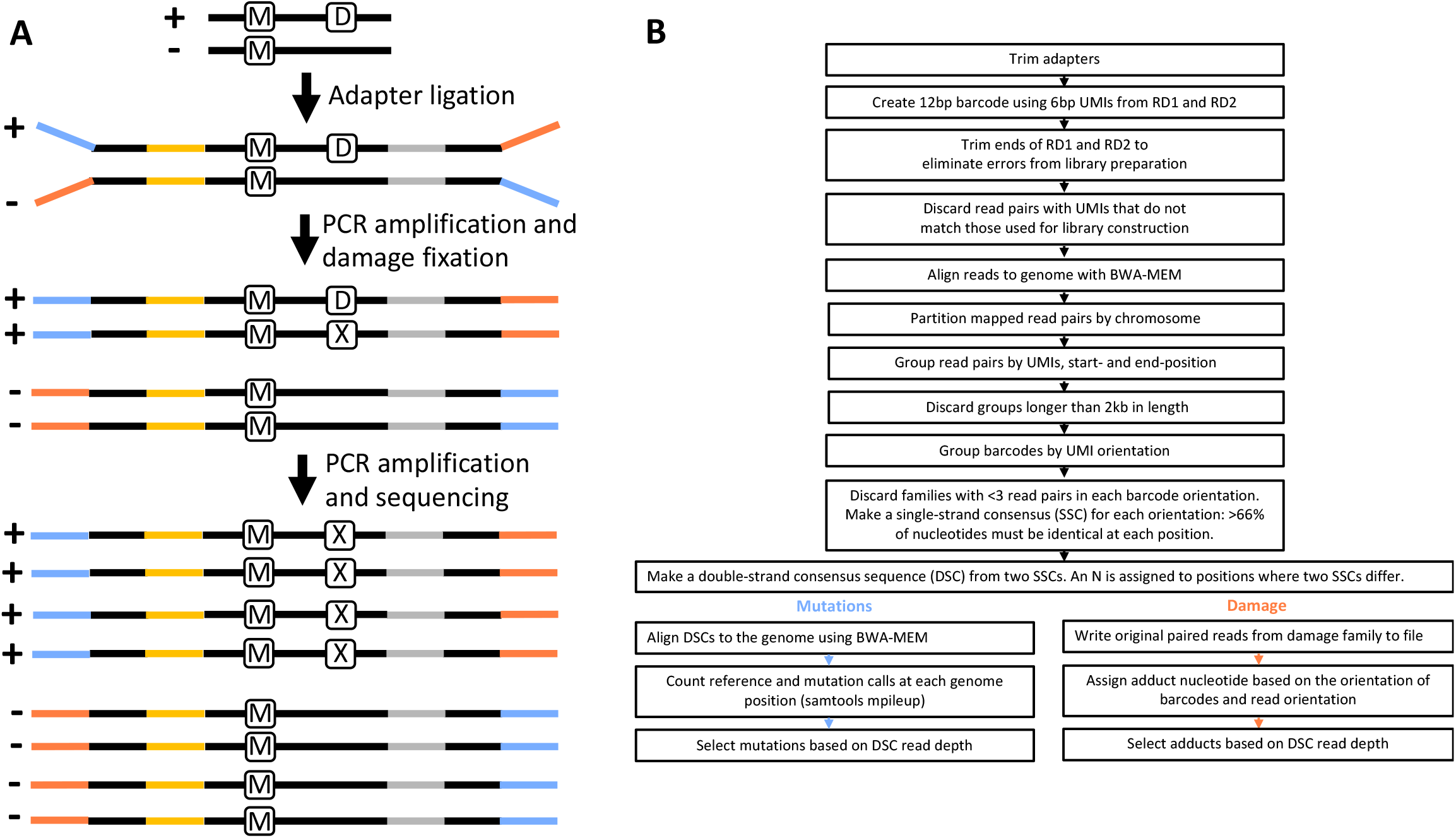
Outline of MADDD-Seq method. **A)** MADDD-Seq distinguishes mutations from artifacts introduced by sequencing technology or NGS-protocols by confirming that a true mutation (M) is present on both strands of a DNA duplex, while artifacts are only present in one. In addition, MADDD-Seq can identify damaged bases by virtue of the *ex vivo* mutations that the damage induces during the PCR steps required for library preparation (X, for *ex vivo* mutation). These mutations arise exclusively and repeatedly on copies that are made from the damaged strand. The + and – symbols refer to the plus and minus strand of a DNA duplex. Directional adapters are depicted as orange and blue lines. UMI’s are depicted as yellow and grey lines (see text for further explanation). **B)** Bio-informatic workflow of MADD-seq data analysis

This barcoding strategy resembles the forked adapters used in traditional duplex sequencing techniques. However, MADDD-seq adapters carry different sequences on their forked ends, and alternate in their orientation towards the UMIs present on each molecule. For example, in the schematic presented in **figure 1**, the orange adapter is always adjacent to the purple UMI on the top strand but paired with the green UMI on the bottom strand. Thus, molecules that derive from the top strand always read blue fork – yellow UMI – insert – grey UMI – red fork, while molecules from the bottom strand read blue fork – grey UMI – insert – yellow UMI – orange fork. Our bio-informatic pipeline uses this information to distinguish between copies that are made from the top or bottom strand of a DNA duplex, so that DNA damage can be localized to a single base, in a single strand of the original DNA duplex.

## RESULTS

To provide a proof of concept for our strategy, we synthesized a 50bp double-stranded oligonucleotide that carries an O^6^-methyl guanine adduct at the 11^th^ position. O^6^-methyl guanine (O^6^-meG) lesions are small but highly mutagenic adducts that arise due to endogenous and exogenous processes alike(*22*), and are associated with a wide variety of spontaneous and environmental cancers(*22, 23*). In addition, O^6^-meG lesions were recently associated with Alzheimer’s disease(*24*). Thus, a sensitive assay for the detection of O^6^-meG lesions and the mutations they cause is highly desirable and could be an important factor for disease prognosis, tumor grading, and the prediction of treatment efficacy. We found that MADDD-seq correctly identified an O^6^-meG adduct at the 11^th^ position in at least 94% of the sequenced oligonucleotides (**fig. 2a-b**). In 5% of cases, an undamaged base was called, while the damaged base was mistaken for a mutation in 0.89% of cases. The relative inaccuracy of inserting an O^6^-meG base at this position during oligo synthesis may account for most of these undamaged base calls (only >85% of oligos are guaranteed to contain the desired O^6^-meG lesion by GeneLink). Regardless, this data demonstrates that that O^6^-meG lesions allow for efficient error-prone bypass by the DNA polymerases present in our PCR amplification mixture, and that as a result, mutagenic O^6^-meG lesions can be detected with considerable accuracy.

**Figure 2.**
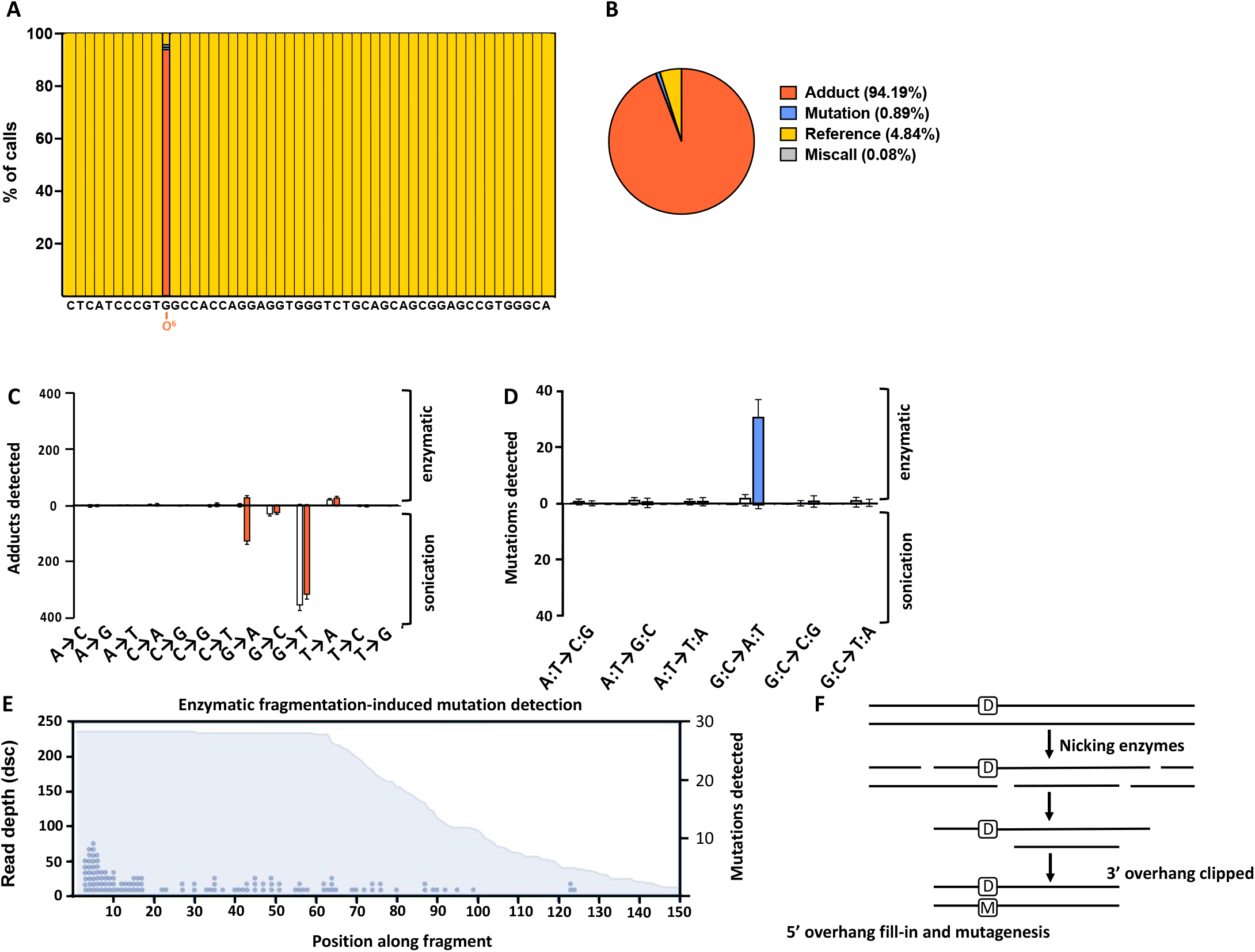
Detection of an O6-meG lesion in a synthetic oligo. **A**. An oligo was synthesized that carries an O^6^-methyl guanine base at the 11^th^ position and annealed to a complementary strand to create a DNA duplex. This oligo was processed with standard MADDD-seq protocols and pipelines to detect both DNA damage and mutations. MADDD-seq correctly identified an O^6^-methyl guanine adduct at the 11^th^ position (orange bars) in nearly all sequenced oligos, while it identified undamaged reference bases at all other positions (yellow bars). **B**. Deeper analysis demonstrated that at the 11^th^ position, MADDD-seq identified an O^6^-methyl guanine adduct in 94% of cases, a undamaged reference base in 5% of cases, and a mutation in 0.89% of remaining cases. For **A** and **B**, O^6^-meG adducts are depicted in orange, mutations in blue, reference bases in yellow and miscalls in grey. **C**. Isolated DNA was either treated with MNNG (orange bars) or not (white bars), and MADDD-seq was used to detect O^6^-meG lesions (G→A transitions). This DNA was either sheared by sonication or enzymatic digest. **D**. Next, we monitored the same isolated DNA samples for mutations. Treated samples are depicted with blue bars, and untreated samples with white bars. In both **C** and **D**, error bars indicate SEM, n=3. **E**. Mutations found in enzymatically fragmented samples are biased toward the 5’end of the fragment, suggesting a non-biological assay-induced explanation for the presence of these mutations. The blue surface depicts the read depth of the fragments we sequenced, while the blue dots indicate the number of mutations we detected. One dot indicates one mutation. **F**. Schematic of how large overhangs generated by enzymatic fragmentation may convert DNA damage (D) into mutations (M) in pre-PCR gDNA libraries.

In biologically relevant scenarios (such as acute exposure to a DNA damaging agent) O^6^-meG adducts are distributed randomly across a complex genome that contains countless structural and functional features and displays an elaborate 3D architecture. To determine if MADDD-seq can identify O^6^-meG adducts under those conditions as well, we treated isolated DNA from the budding yeast *S. cerevisiae* with MNNG, a DNA alkylating agent that induces O^6^-meG lesions. We performed this test on isolated DNA to prevent the treatment from inducing mutations, allowing us to test the ability of MADD-seq to distinguish between actual mutations and damaged bases. MADDD-seq correctly identified a substantial increase in O^6^-meG lesions after MNNG exposure, while no mutations were induced by the treatment. Moreover, MADDD-seq detected these lesions on all 16 chromosomes, as well as the mitochondrial genome, indicating that it can be used to detected O^6^-meG adducts in a true genome-wide fashion. Importantly, MADDD-seq was only capable of distinguishing between DNA damage and mutations if the DNA was sheared by sonication (**fig. 2c-d**). If the DNA was fragmented enzymatically, MADDD-seq reported a substantial increase in mutations instead (**fig. 2e**). These mutations were biased toward the end of sequenced DNA fragments, suggesting a non-biological explanation. We suspect that these artifacts are created by overhangs and gaps that are generated by the enzymatic fragmentation mix. Potentially, these single-stranded regions are then filled in by DNA polymerases included in the enzymatic mixture to create mutations opposite O^6^-meG bases (**fig. 2f**). Thus, the DNA fragmentation method is essential to the success of MADDD-seq, suggesting that it may be important for other duplex sequencing techniques as well.

After establishing the optimal protocol to detect DNA damage with MADDD-seq, we wanted to test its ability to detect DNA damage and mutations simultaneously, with a single assay. In addition, we wanted to perform this experiment in a biologically relevant scenario, by mimicking an acute exposure to a mutagenic compound (**fig. 3a**). To do so, we exposed rapidly dividing cells of the budding yeast *S. cerevisiae* to MNNG for 40 minutes, washed the cells to remove MNNG from the medium and then tracked both adducts and mutations over a 24-hour timespan. As expected, MADDD-seq identified a large increase in O^6^-meG lesions immediately after exposure, similar to the experiments described above. These lesions covered the entire genome in a relatively unbiased fashion and affected protein coding regions, intragenic sequences, introns, untranslated regions, and regulatory elements alike (**fig. 3b-c**). We found that on average, yeast cells carried approximately 550 O^6^-meG lesions per million bases (Mb) after exposure, which amounts to approximately 1 lesion every 2,000 bases. Moreover, these lesions rapidly declined over a 24-hour time span (**fig. 3d**), which we ascribed to the activity of both DNA replication and the DNA repair protein MGT1, the yeast homologue of MGMT(*25*). We then analyzed our dataset with the second arm of our bioinformatic pipeline specifically designed to detect mutations.

**Figure 3.**
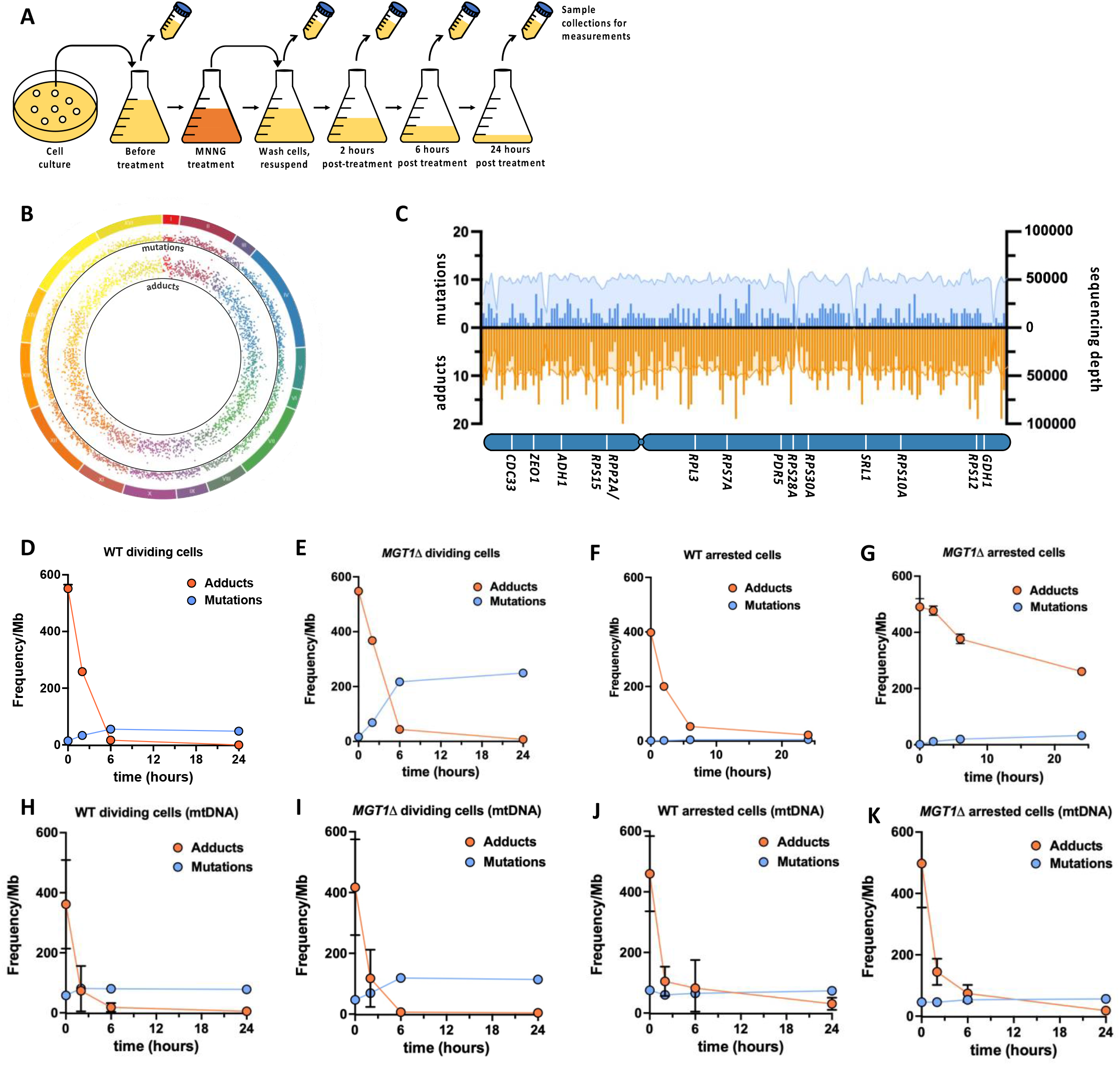
Simultaneous detection of DNA damage and mutations in yeast cells withg MADDD-seq. **A**. Schematic representation of experimental procedures. Single colonies were inoculated in liquid medium and expanded into a 250ml culture. Over the course of 24 hours, multiple 50ml aliquots of cells (conical tubes) were collected at different timepoints to detect DNA damage and mutagenesis by MADDD-seq. **B**. Circos plot of all mutations and adducts detected in this study. The outer ring depicts the chromosome and the location along the chromosome that mutations and adducts were detected at. Each dot in the inner ring depicts the number of adducts detected in 5,000bp intervals. The further away from the center of the plot, the more adducts were detected. The middle ring depicts the number of mutations detected in the same 5,000bp intervals. The further away from the center of the plot, the more mutations were detected. **C**. MADDD-seq coverage and endpoint detection across chromosome XV. Coverage is depicted by the colored surface, while mutations are depicted as orange bars, and adducts as and blue bars, binned in 5,000bp intervals. This plot contains all the mutations and adducts detected in this study. **D-G**. Presence of DNA mutations (blue lines) and adducts (orange lines) over a 24-hour timespan in the nuclear genome of dividing and arrested cells, with or without deletion of MGT1. **H-K**. Presence of DNA mutations (blue lines) and adducts (orange lines) over a 24-hour timespan in the mitochondrial genome of dividing and arrested cells, with or without deletion of MGT1. For D-K, error bars indicate upper and lower limits of confidence intervals for n=1.

Consistent with the mutagenic nature of O^6^-meG, we found the mutation frequency rapidly increased over time (**fig. 3d**), reaching 48 mutations per Mb after 24 hours. Because 550 O^6^-meG lesions per Mb were induced after exposure, roughly 9% of O^6^-meG lesions were fixed into mutations, while 91% were repaired. This straightforward calculation demonstrates how simultaneous detection of DNA damage and mutations allows mutagenic endpoints to be quantified in a direct, highly informative manner.

Next, we wanted to test the ability of MADDD-seq to detect DNA damage and mutations in a medically relevant scenario. To do so, we mimicked MGMT hypermethylation in brain tumors by deleting *MGT1* in yeast cells and treating these cells with MNNG to induce O^6^-meG lesions. Immediately after exposure, *Mgt1Δ* cells and WT cells carried the same number of O^6^-meG lesions, indicating that the presence or absence of MGT1 does not affect the total number of lesions that arise after acute exposure. However, *Mgt1Δ* cells converted these lesions into mutations at a much higher rate, increasing the mutation frequency (**fig. 3e**) to 249 mutations per Mb, demonstrating that 45% of the O^6^-meG lesions were fixed into mutations. These observations underscore the importance of MGT1 for the repair of O^6^-meG adducts and illustrate how MADDD-seq could be deployed in biomedical research or the clinic to investigate the consequences of genetic and epigenetic alterations to MGMT in human tumors.

In addition to rapidly dividing cells (like the cells present in a brain tumor), non-dividing cells can become deficient for MGMT as well. For example, it was recently shown that female Alzheimer patients display hypermethylation of the MGMT promoter, leading to reduced MGMT expression, while males do not(*24*). We recently confirmed this observation in a separate cohort of patients (Chang, BioRxiv 2023). To mimic MGMT hypermethylation in neurons and explore the impact of mutagen exposure on non-dividing cells, we arrested WT and *Mgt1Δ* cells with α-mating factor and treated them with MNNG according to the protocol described above. In WT cells, we found that 400 O^6^-meG lesions per Mb were present immediately after exposure, and as in dividing cells, these lesions were rapidly removed from the genome (**fig. 3f**), indicating that the proliferative state of the cells does not affect the rate with which O^6^-meG lesions are removed from the genome. In contrast though, we found that the mutation rate was profoundly affected by the proliferative state. Despite extensive damage, the mutation frequency only rose to 5 mutations per Mb over a 24-hour timespan (compared to 48 in dividing cells), suggesting that only 1.2% of these lesions were converted into mutations. It should be noted though, that after multiple hours of arrest, some cells escaped arrest by α-mating factor and lost their arrested state. It is likely that the mutations we detected in the arrested cells arose in these newly dividing cells. These observations suggest that cell division and DNA replication are an important step for O^6^-meG lesions to be fixed into mutations, an observation what was previously made for other DNA lesions as well, including cyclobutane pyrimidine dimers(*26*). Ref bielas papers As expected, we further found that arrested *Mgt1Δ* cells retained O^6^-meG lesions for an extended period of time (**fig. 3g**). Even after 24 hours, 53% of lesions remained, compared to only 5% in WT cells. This observation underscores the importance of MGT1 for the repair of O^6^-meG lesions in non-dividing cells and suggests that in non-dividing cells that lack MGMT, O^6^-meG lesions are retained for an extended period of time. Finally, we observed a limited number of mutations in arrested *Mgt1Δ* cells (**fig. 3g**), which most likely arose in cells that escaped α-mating factor arrest, similar to WT cells.

One exciting aspect of MADDD-seq is that it can detect mutations and DNA damage in a true genome-wide fashion, which includes mitochondrial DNA (mtDNA). Thus, the mutation rate and the kinetics of DNA repair can be directly compared between nuclear and mtDNA. Intriguingly this comparison indicated that although the mitochondrial genome is extensively damaged after MNNG exposure (although less so than the nuclear genome), mutations rarely arose (**fig. 3h-k**). We previously made similar observations with other chemicals and forms of DNA damage(*27*), indicating that mtDNA is highly resistant to mutation accumulation due to DNA damage. Although we did detect a clear increase in mtDNA mutations in cells that lacked MGT1, this increase was unusually modest compared to the nuclear genome. In addition, we found that even in the absence of MGT1, the mitochondrial genome is almost completely cleared of DNA damage over a 24 hour timespan (**fig. 3h-k**). Thus, even though the mitochondrial genome has a higher somatic mutation rate when compared to the nuclear genome, and is thought to lack a comprehensive DNA repair system(*28*), mutagen-induced mutations rarely arise, and lesions do not persist for long periods of time. It will be interesting to dissect the mechanisms by which mitochondria achieve this remarkable feat.

The experiments described above demonstrate that MADDD-seq can identify mutations and DNA damage across the genome in a massively parallel fashion. As a result, we hypothesized that MADDD-seq datasets can be mined for known and unknown variables that alter the sensitivity of DNA sequences to mutagens, the pathways that are used for DNA repair, or the propensity for damage to be fixed into mutations. To test this hypothesis, we first decided to investigate the mechanism that is responsible for the limited repair of O^6^-meG lesions in the absence of MGT1. Interestingly, we found that in *Mgt1Δ* cells, repair of O^6^-meG lesions is more pronounced on the transcribed strand of protein coding genes compared to the non-transcribed strand, a distinction that is not visible in WT cells (**fig. 4a-b**). This observation strongly suggests that in the absence of MGT1, transcription-coupled DNA repair plays an important role in removing O^6^-meG lesions from the genome. To explore this possibility further, we generated custom-made transcriptome datasets of treated and untreated yeast cells in a dividing or arrested state and found that genes with higher expression levels indeed displayed faster DNA repair, specifically on the transcribed strand. Again, no such distinction was visible in WT cells (**fig. 4c-f**). These observations demonstrate that MADDD-seq can be used to identify DNA repair pathways responsible for the clearance of DNA adducts from the genome.

**Figure 4.**
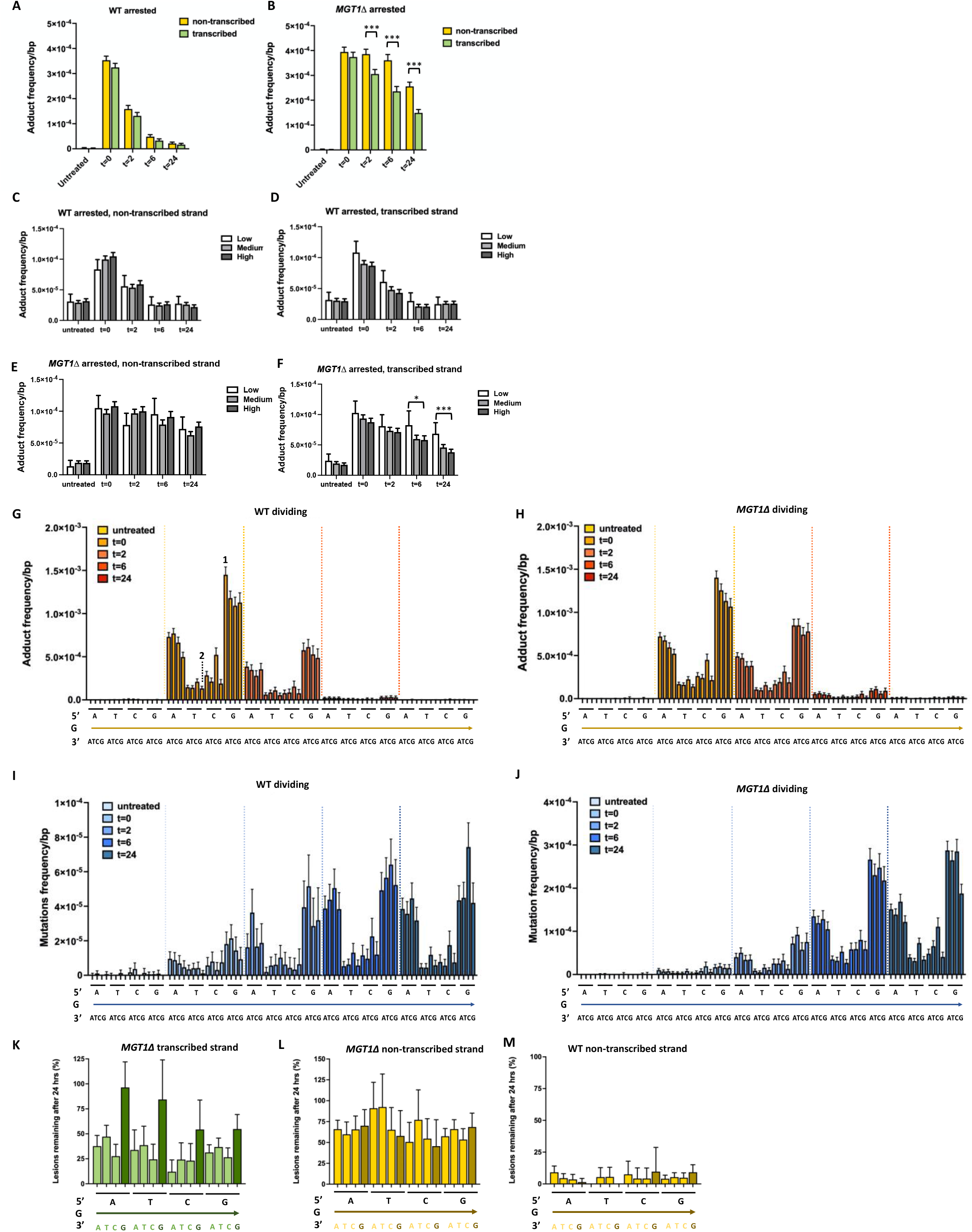
Adduct repair and mutations induction as a function of genetic context and transcription level. **A-B**. O^6^-meG lesions are removed faster from the transcribed strand compared to the non-transcribed strand in arrested *Mgt1Δ* cells, but not in WT cells. **C-F**. Based on transcriptome analysis (n=6 for all conditions), genes were divided into 3 bins: low, medium and high transcription level and the frequency of O^6^-meG lesions was monitored across these three bins over a 24-hour timespan. We found that O^6^-meG lesions were removed faster from highly transcribed genes compared to rarely transcribed genes, but only in the transcribed strand of arrested *Mgt1Δ* cells. **G-J**. The frequency of O^6^-meG lesions (orange bars) and mutations (blue bars) were monitored over a 24-hour timespan as a function of the genetic context in which the damaged base was present. The base that flanks the O^6^-meG lesion on the 5’ side is depicted immediately underneath the graph. The damaged guanine base itself is depicted underneath the 5’ base (and is present in all triplets, as depicted by the arrow), while the base flanking the O^6^-meG lesion on the 3’ side is depicted below the arrow. For example, at t=0, O^6^-meG lesions are most common at guanine bases that are flanked on its 5’ side by another guanine, and on its 3’ side by adenine (labeled by the number 1). In contrast, guanine bases that are flanked on its 5’ side by thymine and on its 3’ side by guanine are the least likely to carry an O^6^-meG lesion (labeled by the number 2). For comparisons between adducts and lesions, please compare the adducts at t=0 (when the maximum number of lesions are present) to the mutations at t=24 (when all these lesions have been fixed into mutations). **K**. The transcription-coupled DNA repair process that removes O^6^-meG lesions from the genome prefers a genetic context in which a guanine base (dark green) is not present on the 3’ side of the damaged base. This preference is not observed in the non-transcribed strand of *Mgt1Δ* cells (**L**), or the transcribed strands of WT cells (**M**). For **L** and **M**, the 3’ guanine base is labeled in dark yellow. Statistical tests for significance were performed using tests of equal proportion with Yates continuity correction. * = p<0.05, ** = p<0.01, *** = p<0.001. For all conditions, n=1.

In addition, we decided to use our MADDD-seq dataset to identify new variables that link DNA damage to mutations. For example, mutations are frequently found to arise in a non-random fashion after mutagen exposure, with the rate of mutation changing as a result of genetic context(*29*). Our data displays a similar trend, with mutations arising more frequently on damaged guanine bases flanked by a purine on their 5’ side compared to a pyrimidine (**fig. 4g-j**). To shed light on the mechanism responsible for this observation, we examined the distribution of O^6^-meG lesions across the genome and found that they preferentially arise on guanine bases flanked by 5’ side purines as well (**fig. 4g-h**). Thus, this observation suggests that the genetic context in which mutations arise is not dictated by a preference of DNA repair or the mutagenic process itself to a specific genetic context, but is primarily caused by the specificity with which the O^6^-meG lesions accumulate on the genome. We did find evidence though, that the transcription-coupled DNA repair process that takes over in the absence of MGT1 may contribute the mutagenic properties of O^6^-meG to a limited degree. Most notably, we found that O^6^-meG lesions that are flanked on their 3’ side by a second guanine are repaired less efficiently compared to adenine, cytosine or thymine (**fig. 4k**). We did not observe this discrepancy on the non-transcribed strand of *Mgt1Δ* cells (**fig. 4l**) or in the transcribed strand of WT cells (**fig. 4m**), indicating that it is specific to the transcription coupled DNA repair mechanism active in *Mgt1Δ* cells. Taken together, these observations demonstrate how MADDD-seq datasets can be used to come to make highly detailed observations that explore key aspects of the relationship between DNA damage, DNA repair and mutagenesis.

## DISCUSSION

Mutations play an important role in human aging and disease. However, because they tend to be rare, and are distributed randomly across in the genome, it can be difficult to detect them among the overwhelming background of WT DNA. To solve this problem, a number of highly sophisticated genome-wide detection techniques have been developed that greatly improve the detection of mutations, including Cypher-seq(*13*), circle-sequencing(*30*), o2n-sequencing(*31*), SMM-seq(*32*) and duplex sequencing(*19*).

Although DNA damage is more abundant than mutations, it has proven equally difficult to detect. In addition to the problems highlighted above, a third complication is that DNA damage comes in many shapes and sizes, with each lesion representing a new challenge to overcome(*12*). Despite these challenges, multiple highly advanced and often complementary techniques have been developed over the past decades. For example, mass spectrometry(*33*) and radioactive labeling techniques(*34*) now allow researchers to determine the total amount of DNA damage present in a biological sample, while the precise locations of lesions can be identified with lesion-specific sequencing techniques using antibodies and modified enzymes(*5, 7-9, 35-46*) or long-range PCR assays(*47-50*). It is important to note that the sensitivity of these techniques is highest when they target bulky lesions, while smaller lesions tend to be missed. For example, antibodies raised against DNA adducts work best when they target bulky lesions that are easy to discriminate from undamaged bases, while long range PCR reactions require lesions to block DNA polymerases during DNA synthesis, which is more likely to happen in the case of bulky adducts(*27*). In contrast, small lesions are difficult to discriminate from undamaged bases and allow for more efficient translesion synthesis during DNA synthesis. It is precisely for these reasons though, that these lesions are highly mutagenic and thus dangerous for human health. To address this issue, we developed MADDD-seq, an assay that is specifically designed to detect small lesions that are highly mutagenic in nature.

In the proof-of-principle experiments described here, we demonstrate that MADDD-seq can efficiently distinguish between O^6^-meG lesions and mutations in DNA samples extracted from eukaryotic cells and do so in a genome-wide fashion. Importantly, O^6^-meG lesions are associated with a wide variety of human cancers. For example, the DNA repair protein O^6^-methylguanine-DNA methyltransferase (MGMT) is hypermethylated in 50% of grade IV glioblastomas, 85% of thyroid cancers and 70% of colorectal cancers(*51, 52*), suggesting that O^6^-meG lesions are the primary source of the mutations that drive these cancers. In addition, MGMT is hyper methylated in female (but not male) patients with Alzheimer’s disease(*24*), which could contribute to the increased mutagenesis and apoptosis seen in hippocampal neurons of patients (*53*), as well as the sexual dimorphism of the disease itself. Thus, a sensitive assay for the detection of O^6^-meG lesions and the mutations they cause is highly desirable and could be an important factor for disease prognosis, tumor grading, and the prediction of treatment efficacy. Here, we demonstrate that MADDD-seq data can be analyzed to provide precise insight into the distribution of DNA lesions in both nuclear and mitochondrial DNA, the kinetics of DNA repair across these genomes, and the rate with which lesions are fixed into mutations. This data can then be parsed through multiple filters to determine the structural, functional or sequence specific parameters that control these processes. For example, by treating WT and *Mgt1Δ* cells yeast cells with MNNG, MADDD-seq clearly outlined the distribution of O^6^-meG lesions across the genome, demonstrated the kinetics of DNA repair and mutagenesis in WT and *Mgt1Δ* cells and allowed us to quantify how many mutations arose as a result of exposure in both nuclear and mitochondrial DNA. Remarkably, these observations ran counter to the idea that mtDNA is more prone to mutagenesis due to its reduced DNA repair capacity, as lesions were rapidly cleared, and only a limited number of mutations arose. In addition, we were able to parse our datasets to demonstrate that transcription coupled DNA repair provides a back-up mechanism for the repair of O^6^-meG lesions in yeast cells, that this DNA repair pathway prefers certain genetic contexts, and that the distribution of mutations after exposure is primarily dictated by the initial distribution of lesions across the genome.

It is further important to note that even though we performed our experiments with MNNG, and monitored O^6^-meG lesions after MNNG exposure to provide a proof of principle for our approach, it is likely that MADDD-seq can be used to detect other lesions as well. As long as a lesion allows for efficient translesion synthesis and is mutagenic in nature (such as 8-oxo-guanine and uracil), it should be possible to detect it with MADDD-seq. It may even be possible to identify multiple lesions at once, based on the mutation spectrum they produce. For end users that want to adapt MADDD-seq for their lesion of interest, we highly recommend using an analog approach to the one described above.

Another advantage for end users is that MADDD-seq is allows mutations and DNA damage to be detected simultaneously, with a single assay, using a single sample in a single reaction. Thus, DNA damage and mutations are always detected under identical conditions, thereby limiting unintended artifacts due to sample handling, batch effects or user errors. Moreover, having a single assay for multiple endpoints means that precious material such as human brain samples or tumor biopsies can be preserved for additional experiments in the future. In addition, it cuts back on the time and money needed to perform experiments, the expertise required to run multiple assays and the need to troubleshoot multiple complex techniques when they provide contradictory results. In doing so, we expect that MADDD-seq will prove to be a versatile, multi-functional detection tool for mutation research that can help basic researchers as well as clinical scientists reveal new biology that underlies human aging and disease.

## METHODS

### MADDD-Seq Assay (Mutation and DNA Damage Detection-Sequencing)

#### Concept

gDNA fragments containing adducts and/or mutations are ligated to custom MADDD-Seq barcoded adapters. The adapters contain one of 218 known 6-bp unique molecular index (UMI) and a forked tail of two non-complementary sequences (blue and orange in fig. 1). A T-overhang on the UMI end of the adapter can be ligated to either end of an a-tailed gDNA fragment. The random pairing of two UMI’s ligated to the same fragment (purple and green in fig. 1) in combination with the fragment’s location on the genome are used to identify each unique gDNA fragment. Amplification of this starting fragment generates multiple PCR duplicates. Through their position in the genome, UMI1, and UMI2, we can identify ‘families’ of PCR duplicates that all originated from the same unique starting molecule. Mutations present in the original molecule will be present in most if not all PCR duplicates. However, errors arising from PCR amplification or next-generation sequencing will not be found across a family of PCR duplicates. Therefore, we can distinguish a true mutation, present in the unique starting gDNA fragment, from NGS and PCR-induced errors.

#### gDNA Fragmentation

Initial library preparation utilized the NEBNext ^®^ Ultra™ II FS Enzyme Mix and NEBNext ^®^ Ultra™ II FS Reaction Buffer to fragment, end-repair, and A-tail. However, sonication was implemented to fragment DNA after results suggested enzymatically shearing DNA induced the conversion of single-stranded DNA damage to mutation (fig. **X**) In the sonication protocol, 500 ng of genomic DNA in 50 uL was fragmented to ∼300 bp using Covaris S220 (Peak Power = 175, Duty Factor = 10.0, Cycles/Burst = 200, Run Time = 50s). Fragment sizes were verified using Agilent TapeStation 4150 using D1000 or D5000-HS reagents.

#### Library Preparation: End Repair and Adapter Ligation

The fragmented DNA was end-repaired and A-tailed using the NEBNext ^®^ Ultra™ II End Prep system. The entire pool of fragmented DNA (∼500 ng) was combined with 7 uL NEBNext Ultra II End Prep Reaction Buffer and 3 uL NEBNext ^®^ Ultra™ II End Prep Enzyme Mix and brought to a total reaction volume of 60 uL with TE. The DNA was incubated with the end repair mix was incubated at 20ºC for 30 minutes followed immediately by a Beckmann-Coulter AMPureXP bead cleaning at 1.8X to halt enzymatic activity and eluted in 35 uL of 0.1X TE. The NEBNext Ultra II End Prep protocol suggests a 65ºC heat inactivation step, which we replaced with this bead cleaning step to minimize heat damage on the libraries.

After the DNA had undergone end repair, we ligated our double-stranded barcoded adapters to the library (see **X** for a detailed adapter preparation protocol). The 35 uL of end repaired DNA was combined with 30 uL of NEBNext Ultra II Ligation Master Mix, 1 uL of NEBNext Ligation Enhancer, and 2.5 uL of 15 uM adapters for a total reaction volume of 68.5 uL. This adapter ligation mix was incubated at 20C for 15 minutes before proceeding directly with a Beckmann-Coulter AMPureXP bead cleaning at a concentration of 0.8X to size exclude surplus adapters and eluting in 20 uL 0.1XTE.

#### Library Quantification

The adapter-ligated libraries were quantified by mass using the Invitrogen Qubit 4 Fluorometer with Qubit 1X dsDNA HS Assay Kit reagents. The fragment size and concentration of each sample was quantified using Agilent’s 4150 TapeStation System with D1000 reagents. To get an accurate estimate of the number of adapter-ligated fragments, droplet digital PCR (ddPCR) was performed with primers specific to the u-loop adapters (forward 5’-GACTGGAGTTCAGACGTGTGC-3’ and reverse 5’-CACTCTTTCCCTACACGACGC-3’). The following reagents were combined to form 22.8 μL of template-free master mix for each reaction well: 12 μL QX200 ddPCR EvaGreen® Supermix, 2.4 μL 0.5 μM forward and reverse primers, 8.4 μL NFW, and 1.2 μL of a 10-fold template dilution series from 1:10 to 1:10^6^. Droplets were generated with Droplet Generation Oil for EvaGreen® using the Bio-Rad QX200 Droplet Generator and amplified with the following cycling parameters: denaturation at 95ºC for 5 minutes followed by 40 cycles of a 30 second denaturation at 98ºC and extension at 65ºC for 1 minute. This was followed by 5 minutes at 4ºC then a final step of 5 minutes at 90ºC before cooling again at 4ºC. Droplets were analyzed by the Bio-Rad QX100 Droplet Reader.

#### Amplification for Illumina sequencing

Based on quantification by ddPCR, 3 million molecules were amplified using indexed primers for multiplexing samples during sequencing with an adaptation of NEBNext’s Multiplex Oligos for Illumina universal forward primer (5’-AAT GAT ACG GCG ACC ACC GAG ATC TAC ACT CTT TCC CTA CAC GAC GC-3’) and indexed reverse primers, 5’-CAA GCA GAA GAC GGC ATA CGA GAT **XXX XXX** GTG ACT GGA GTT CAG ACG TGT GC-3’), where the bases ‘**XXX XXX**’ correspond to indices 1,2,4,5,6,7,12,15,16, and 19. Index primers were selected from a combination of Index Primers Set 1 and Set 2 based on the NEB index pooling guide. The primers used are identical to NEBNext’s Multiplex Oligos for Illumina except for the removal of 10 bp from the 3’ end of each primer complementary to the double-stranded portion of the adapters to prevent these primers from annealing with molecules from the opposite strand and conflating the sequence-based strand distinction in our assay.

The amplification reaction contained 25 μL NEBNext Q5 2X MM, 2.5 μL 10 μM NEBNext Universal Primer, 2.5 μL 10 μM NEBNext Index Primer, an aliquot of 3 million copies of template, and NFW to bring the total volume to 50 μL. Cycling parameters for amplification were as follows: initial denaturation at 98ºC for 30 seconds, followed by 15 cycles of denaturation at 98ºC for 10 seconds and extension at 65ºC for 1 minute. A final step of 65ºC for 5 minutes was performed prior to sample cooling at 4ºC. Two successive AMPure XP bead purification of PCR products were performed at 0.7X concentration, and the samples were eluted in a final volume of 20 μL 0.1X TE.

#### Amplified library quantification and re-amplification

Amplified libraries were quantified by Invitrogen’s Qubit 4 Fluorometer with Qubit 1X dsDNA HS Assay Kit reagents and Agilent’s 4150 TapeStation System with D5000 High Sensitivity reagents. ddPCR was performed using primers complementary to NEBNext’s Multiplex Oligos for Illumina: forward 5’-AAT GAT ACG GCG ACC ACC GA-3’ reverse 5’-CAA GCA GAA GAC GGC ATA CGA-3’ and a FAM fluorescent probe complementary to the Illumina primers to specifically measure the number of DNA fragments that could be sequenced (5’-/56-FAM/CCC TAC ACG /ZEN/ACG CTC TTC CGA TCT/3IABkFQ/-3’). The reaction mixture contained 12 μL ddPCR Master Mix for Probes, 3 μL 2 μM DPL probe, 2.16 μL 10 μM primers, 1.2 μL template along a 1:10 – 1:10^6^ dilution gradient, and NFW to 24 μL. After droplet generation using the QX200 droplet generator and ddPCR Droplet Generation Oil for Probes (Bio Rad), reactions were heated at 95ºC for 10 minutes, then denatured at 95ºC for 30 seconds and extended at 65ºC for 1 minute for 40 cycles before heating at 98ºC for 10 minutes and cooling at 4ºC. Droplets were analyzed on the Bio Rad QX100 Droplet Reader.

Based on ddPCR quantification, samples that were not concentrated enough to sequence were re-amplified to target ideal sequencing concentration. The remaining sample volume was amplified in a 50 μL reaction mixture along with 2 μL NEBNext Q5 2X MM, 5 μL 10 μM library primers (forward 5’-AAT GAT ACG GCG ACC ACC GA-3’ reverse 5’-CAA GCA GAA GAC GGC ATA CGA-3’), and NFW to 50 μL. Cycling parameters for amplification were as follows: initial denaturation at 98ºC for 30 seconds, followed by a variable number of amplification cycles of denaturation at 98ºC for 10 seconds and extension at 65ºC for 1 minute targeting a final concentration of 1.5 nM. A final step of 65ºC for 5 minutes was performed prior to sample cooling at 4ºC. Libraries were purified using 0.9X AMPure XP bead cleaning and eluted in 15 μL 0.1X TE. Library quantification was reassessed using TapeStation, Qubit, and ddPCR for Probes as described directly above. Re-amplification and quantification were repeated as necessary until libraries reached approximately ∼0.2-1.0 nM. Target concentrations are optimized to be the lowest possible given the total quantity of DNA in order to sequence as much of the library as possible. Samples were then multiplexed and run on a MiSeq Nano Kit using custom primers specific to the NEBNext Index primers (Read 1 Primer: 5’-ACA CTC TTT CCC TAC ACG ACG CTC TTC CGA TCT-3’, Index Primer: 5’-GAT CGG AAG AGC ACA CGT CTG AAC TCC AGT CAC - GCC AAT - ATC TCG TAT GCC GTC TTC TGC TTG-3’, Read 2 Primer: 5’-GTG ACT GGA GTT CAG ACG TGT GCT CTT CCG ATC T-3’). Based on MiSeq clustering and read distribution, sample volumes were adjusted for equal distribution of reads, pooled, and sequenced on a NextSeq 2000.

#### UDSBC Adapter Preparation

U-loop double stranded barcode adapters (5’-TGA CTA GAT CGG AAG AGC ACA CGT CTG AAC TCC AGT C **dU** A CAC TCT TTC CCT ACA CGA CGC TCT TCC GAT CTA GTC A-s-T-3’) were annealed for 5 minutes at 95ºC then cooled quickly on ice. To prevent cleavage at sites of damage on gDNA, 24 μL of 15μM annealed u-loop adapters were digested prior to ligation using 2 μL USER enzyme in a 40 μL reaction with 1X CutSmart Buffer for 15 minutes at 37ºC. Digested adapters were bead cleaned using AMPure XP beads at 1.8X concentration and eluted in 16 μL 0.1X TE. Note that pre-digestion of adapters results in a reduction in ligation efficiency and that future library preparations will exclude this step.

#### Yeast Culture

Single colonies were inoculated in YAPD and incubated overnight at 30°C in a rotating wheel. In the morning, the optical density (OD_600nm_) of each culture was measured using Thermo Scientific’s Nanodrop 2000C and cells were re-inoculated at an OD_600_ 0.05-0.1 in 50ml YAPD flasks and incubated in an orbital shaker at 30°C. Cells were then grown to an OD of 0.25-0.5 and either harvested or arrested with 50 μg/ml α-mating factor. After 2.5 hours, the cells were visualized under a microscope to confirm they were arrested, and then treated (or not) for 40 minutes with mutagenic compounds 10μg/ml MNNG. After treatment, the cells were washed 3 times in PBS with α-mating factor to remove mutagenic compounds and re-inoculated in YAPD with α-mating factor to maintain arrest over the recovery period.

## SUPPLEMENTAL FIGURES

**Supplementary Figure 1.**
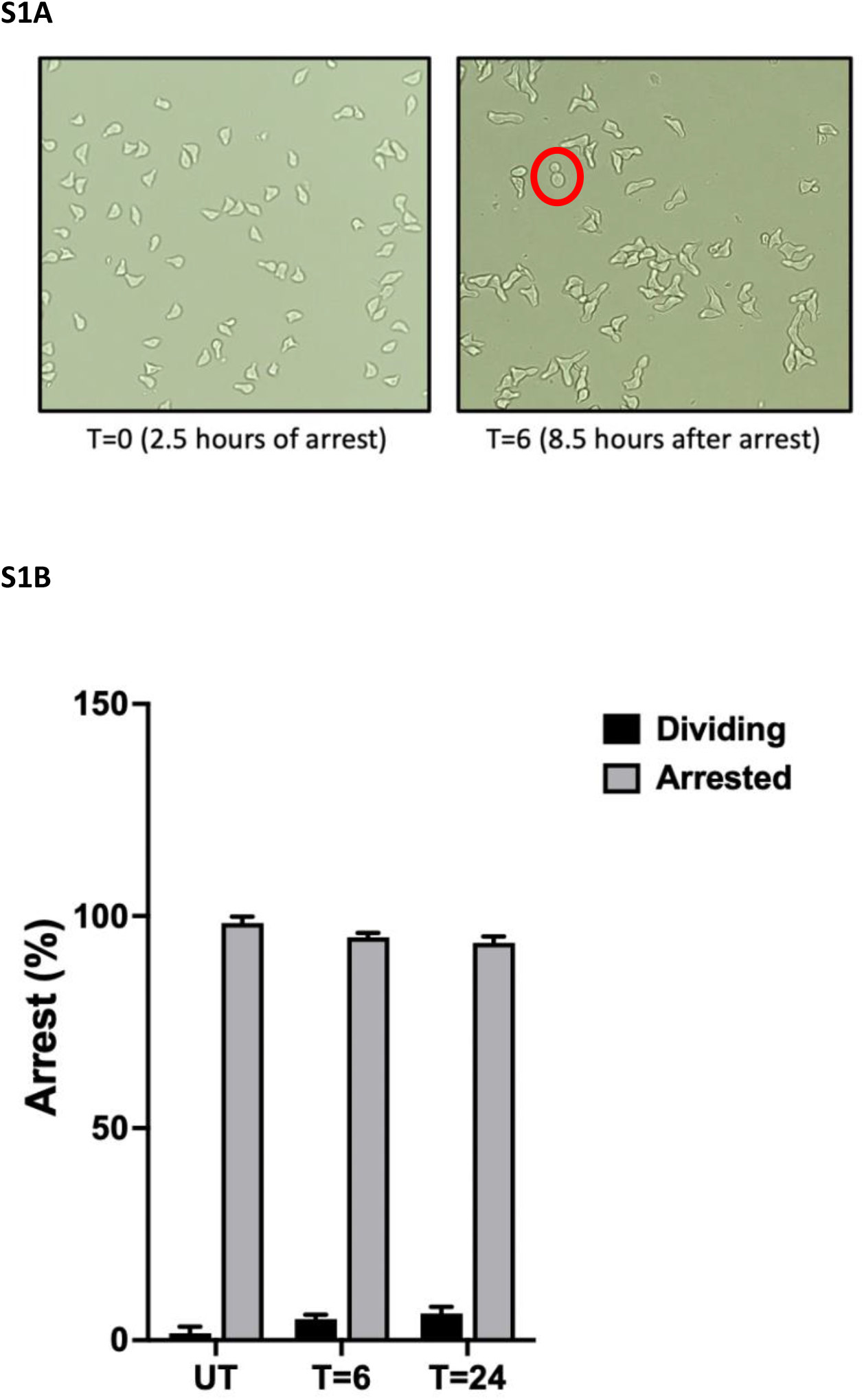
**A**. Cells arrested with α-mating factor were identified by their typical pear-shaped structure (shmooing). In contrast, dividing cells are circular and display a growing bud. One dividing cell, identified by the round mother cell and her growing bud, is depicted inside the red circle. **B**. Cells gradually lose arrest over a 24-hour timespan.

## ACKNOWLEDGEMENTS

We acknowledge support from the National Institute on Aging (R01AG054641 to M.V.), the University of Southern California (SCEHSC pilot award to M.V.), the National Institute of Environmental Health Sciences (U01ES029516 and R01ES026222 to J.H.B.), and the National Cancer Institute (R01CA204894 to J.H.B.).

